# Insights into an alternative pathway for glycerol metabolism in a glycerol kinase deficient *Pseudomonas putida* KT2440

**DOI:** 10.1101/567230

**Authors:** Meg Walsh, William Casey, Shane T. Kenny, Tanja Narancic, Lars M. Blank, Nick Wierckx, Hendrik Ballerstedt, Kevin E O Connor

## Abstract

*Pseudomonas putida* KT2440 is known to metabolise glycerol via glycerol-3-phosphate using glycerol kinase an enzyme previously described as critical for glycerol metabolism (1). However, when glycerol kinase was knocked out in *P. putida* KT2440 it retained the ability to use glycerol as the sole carbon source, albeit with a much-extended lag period and 2 fold lower final biomass compared to the wild type strain. A metabolomic study identified glycerate as a major and the most abundant intermediate in glycerol metabolism in this mutated strain with levels 21-fold higher than wild type. Erythrose-4-phosphate was detected in the mutant strain, but not in the wild type strain. Glyceraldehyde and glycraldehyde-3-phosphate were detected at similar levels in the mutant strain and the wild type. Transcriptomic studies identified 191 genes that were more than 2-fold upregulated in the mutant compared to the wild type and 175 that were down regulated. The genes involved in short chain length fatty acid metabolism were highly upregulated in the mutant strain. The genes encoding 3-hydroxybutyrate dehydrogenase were 5.8-fold upregulated and thus the gene was cloned, expressed and purified to reveal it can act on glyceraldehyde but not glycerol as a substrate.

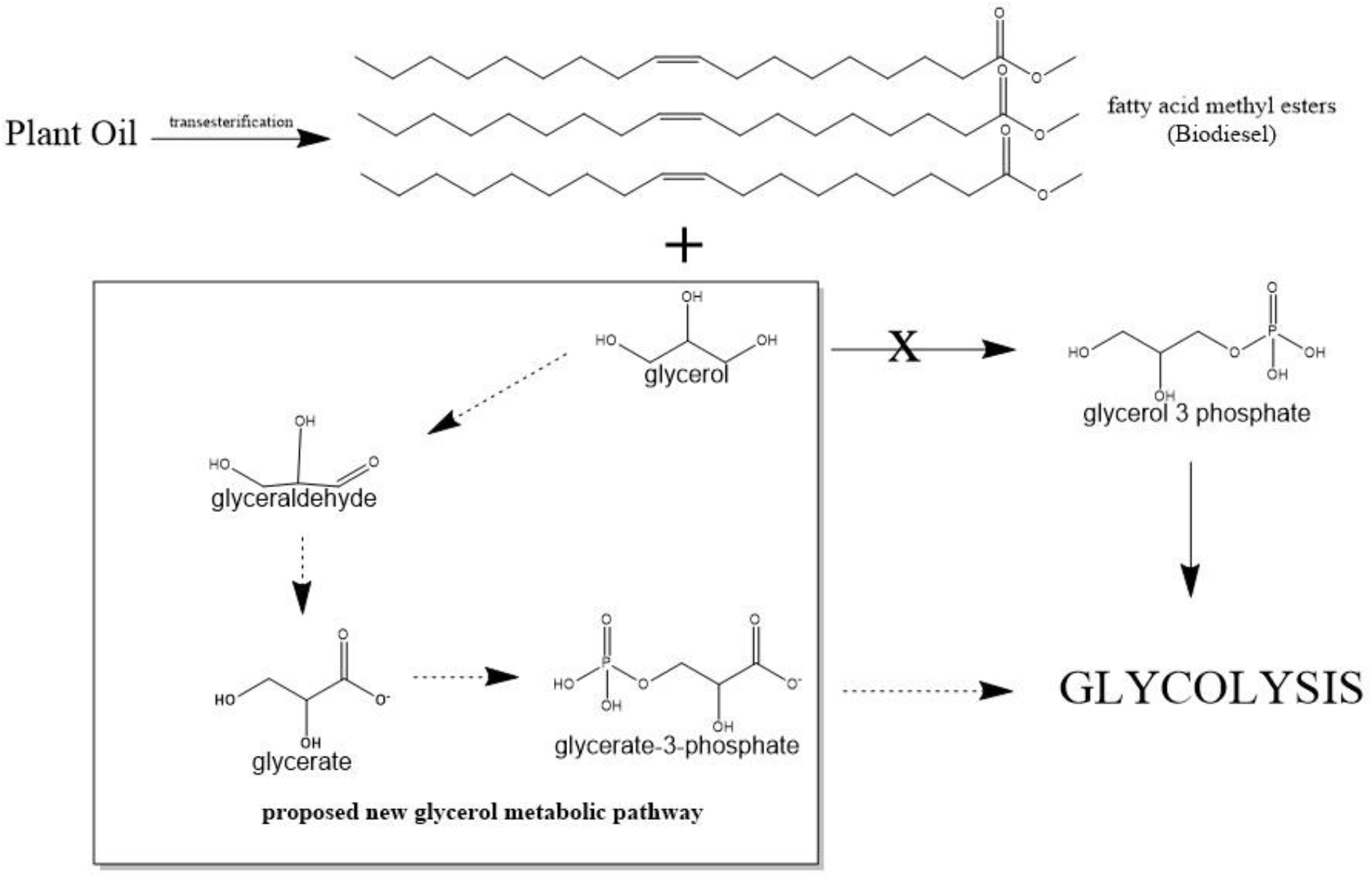

## Introduction

Glycerol is a major by-product of plant oil processing. It is also the major by-product of biodiesel production (10 weight %). An increase in production of biodiesel (2) has led to an increase in the availability of glycerol (3). Glycerol could be an interesting substrate for fermentative production of value added products such as 1,3 propanediol, dihydroxyacetone and polyhydroxyalkanoates (4,5). Polyhydroxyalkanoates (PHA) are polymers that are naturally synthesized and stored in a range of microorganisms including *Pseudomonads* (6). *Pseudomonas putida* KT2440 is a model organism for biocatalysis and synthetic biology. While *P. putida* KT2440 can use glycerol as the sole carbon and energy source for growth, glycerol metabolism in this strain has not been fully characterized. However, the pathway for glycerol metabolism has been extensively studied in the opportunistic pathogen *Pseudomonas aeruginosa* (7–9). As there is high sequence identity in the proposed glycerol metabolic pathway genes in *P. putida* KT2440 with genes in *P. aeruginosa*, some information about glycerol metabolism in *P. putida* KT2440 can be inferred from the research into *P. aeruginosa* (10). In *P. aeruginosa*, the first step in glycerol metabolism is the facilitated diffusion across the cytoplasmic membrane by a glycerol diffusion facilitator (GlpF). The glycerol is then retained intracellularly through phosphorylation by a glycerol kinase (GlpK) to form glycerol-3-phosphate, which cannot diffuse back through the cytoplasmic membrane. In *P. aeruginosa*, the genes for GlpF and GlpK are organised together on one operon (8). The glycerol-3-phosphate is converted to dihydroxyacetone-3-phosphate by a cytoplasmic-membrane associated glycerol-3-phosphate dehydrogenase (GlpD) (11). The Glp operons (GlpFK and GlpD) are negatively regulated by the transcriptional regulator GlpR in *P. aeruginosa* and *P. putida* KT2440 (Schweizer and Po, 1996, Escapa *et al*., 2013). Glycerol-3-phosphate interacts with the GlpR protein, allowing transcription of the glycerol operon. Therefore glycerol-3-phosphate is the true effector of the system. A non-specific kinase must be capable of acting on glycerol or the repression must be leaky because some glycerol-3-phosphate is required to stop repression of the glycerol operon. This repression by GlpR explains the long lag phase when *P. putida* KT2440 is grown on glycerol as the sole carbon and energy source (10).

We constructed a knockout mutant of the glycerol kinase gene (*glpK*). Deletion of this gene severely retarded growth on glycerol as the sole carbon source but did not remove it completely. We therefore undertook transcriptomic and metabolomic studies to understand the metabolism of glycerol in this strain in the absence of glycerol kinase activity.

## Methods

### Strain Manipulation

The Δ*glpK* mutant was constructed by replacing the coding region of the gene with a selective gentamycin resistance cassette as previously described for polyphosphate kinase (*ppk*) gene deletion (12). A complemented mutant strain was produced with an inducible copy of the wild-type *glpK* harboured on a pJB861 expression vector. The resulting *glpK* gene was now under the control of a P_m_ promoter which is stimulated by the XylR protein in the presence of an m-toluic acid inducer.

### Growth and maintenance of strains

*P. putida* KT2440 was maintained on LB agar supplemented with carbenicillin (50 mg/l). *P. putida* KT2440 ΔglpK was maintained on LB agar supplemented with gentamycin (50 mg/ml).

Cultures were grown in 250 ml Erlenmeyer flasks containing 50 ml of Minimal Salt Medium (MSM) (13). For biomass determination, flasks were supplemented with 20 mM substrate and incubated at 30°C, shaking at 200 rpm for 48 h. Cells were then harvested, lyophilised and weighed for cell dry weight determination. For growth curves, flasks were supplemented with 50 mM glycerol and incubated at 30°C, shaking at 200 rpm for 72 h for the wild type cells and 300 h for *Δglpk* cells. Samples were taken periodically and OD_540_ measured in a spectrophotometer to estimate growth.

Strains harbouring an exogenous copy of the *glpK* gene, located on the inducible expression vector pJB861, were used to examine the effect of gene complementation. These strains were grown in the same way as the wild type and *ΔglpK* mutant except for the addition of *m*-toluic acid (as a sodium toluate salt). Transcription is initiated from the *P_m_* promoter due to the interaction of the aromatic *m*-toluic acid with the XylS protein, constitutively expressed by the same plasmid expressing the *glpK* gene.

### Preparation of samples for metabolomics analysis

*P. putida* KT2440 and Δglpk cells were grown as for growth curves until they reached an OD_540_ between 0.5 and 1. 9 ml of 100 *%* methanol was aliquoted into polypropylene tubes, the tubes were weighed and placed in a −50°C ethanol bath. Cells (1.5ml) were added into the tubes and the tubes vortexed. Tubes were weighed again to determine the weight of cells added and then centrifuged at 5000 rpm for 5 minutes at −10°C. The supernatant was discarded and the cell pellets stored at −80°C. To extract metabolites, 1 ml of 80 % (v/v) methanol was added to the pellet and the pellet was redissolved completely. The resuspended cells were transferred to an Eppendorf tube and incubated at 95°C for 5 minutes with vigorous shaking. The tubes were placed on ice for 5 minutes and then centrifuged at 5,000 rpm for 5 minutes at −10°C. Supernatants were transferred to a clean Eppendorf tube and stored at −80°C before analysis.

### Metabolomics procedure

200 μL cell-metabolite methanol extracts were added to Inno-Sil desactivated glass-vials (CS Chromatography Service GmbH, Langerwehe, Germany). Subsequently 50 μL of 20 mg/mL O-methoxyamine hydrochloride in pyridine to prevent degradation of ketone and aldehyde groups during drying process *and* 4 μL of 1 mM m-erythritol (as internal standard) were added to the metabolite extract and intensively mixed on a vortex shaker. Samples were dried for 6 h using a speed vacuum concentrator system (1000 rpm, 30°C) equipped with a freeze trap. Dried samples were resuspended for about 3 min on a vortex in 50 μL freshly prepared O-methoxyamine hydrochloride solution (20 mg/mL) in pyridine. For methoximation (1^st^ step derivatization) of carbonyl moieties into corresponding oximes with o-methoxyamine the samples were incubated at 80°C for 30 minutes with vigorous shaking. After a brief centrifugation at 1000 rpm, the procedure was proceeded by the 2^nd^ derivatization step the silylation of polar functional groups (incl. −COOH, -OH, -NH and −SH) to reduce polarity, increase thermal stability and volatility giving TMS-MOX derivatives of the glycolytic intermediates(14,15). 50 μL of N-methyl-N-trimethylsilyl trifluoroacetamide (MSTFA) were added and intensely mixed for 3 minutes. Subsequently, the samples were incubated at 80°C for additional 30 minutes with vigorous shaking. After brief centrifugation at 1000 rpm the samples were used for immediate GC-MS analysis or stored at 4°C for maximal 3 d.

Samples were analyzed on a TSQ 8000 Triple-Quadrupole MS equipped with PTV-injector (split 1:50; 2μl injection volume) and autosampler (Thermo-Fisher Scientific). Chromatographic separation occurred on a CP-9013 capillary column VF-5ms (Agilent; 30m × 0,25mm × 0,25μm + 10m EZ-guard; oven-program: initial 4 min at 60°C, increase 20°C/min to 320°C, final hold 10 min; scan masses 50-800). The separated compounds were ionized by electron ionization (EI) for reproducible and compound-specific spectra to confirm chemical identities by comparing measured spectra with those of existing spectral libraries (e.g., Golm metabolome database (16)) and standards. Generated ms-spectra were analyzed using Xcalibur and AMDIS to differentiate and identify unknown / well-known metabolites and sort co-eluting peaks. Chemical structures of compounds without reliable standards and retention times were analyzed and identified using the commercial NIST-database.

### Preparation of RNA samples for transcriptomic analysis

*P. putida* KT2440 wild type and Δglpk cells were grown as for growth determination and harvested for RNA extraction at mid-log growth phase (approx. OD_540_ of 1). RNA was extracted using a GeneJET RNA purification kit (Thermo Scientific, Dublin, Ireland) according to the manufacturer’s instructions. Samples were sent in duplicate to Baseclear (Leiden, Netherlands) for transcriptomic analysis.

### Transcriptomics procedure

This procedure was performed at Baseclear (Leiden, Netherlands). Ribosomal RNA molecules were depleted from bacterial total RNA using the Epicentre bacteria Ribo-zero rRNA depletion kit. The dUTP method was used to generate strand-specific mRNA-seq libraries (17,18). The Illumina TruSeq stranded RNA-seq library preparation kit was used. The mRNA was fragmented and converted to double-stranded cDNA. DNA adapters with sample-specific barcodes were ligated and a PCR amplification performed. The library was size-selected using magnetic beads, resulting in libraries with insert sizes in the rate of 100-400 bp. The libraries were diluted, clustered and sequenced on an Illumina HiSeq 2500 instrument. The data produced was processed by removing the sequence reads that are of too low quality and demultiplexing based on sample specific barcodes. An additional filtering step was performed using in house scripts to remove reads containing adaptor sequences of Illumina PhiX control sequences.

### Bioinformatics: transcriptome analysis

Transcriptome analysis was also performed at Baseclear. Sequence reads were additionally filtered and trimmed based on Phred quality scores. The filtered/trimmed reads were aligned against the reference sequence AE015451.2 (*Pseudomonas putida* KT2440) using the CLCbio “RNA-Seq” software. Normalised expression values were calculated and compared between the samples. P-values were determined to assign the significance of expression differences between samples. RPKM was the expression measure used. This is defined as the reads per kilobase of exon model per million mapped reads (19). It seeks to normalize for the difference in number of mapped reads between samples as well as transcript length. It is given by dividing the total number of exon reads by the number of mapped reads (in millions) times the exon length (in kilobases). Statistical analysis was performed using Baggerly et al’s Beta-binomial test (20). It compared the proportions of counts in a group of samples against those of another group of samples and is suited to cases were replicates are available in the groups. The samples were given different weights depending on their sizes (total counts). The weights are obtained by assuming a beta distribution on the proportions in a group, and estimating these, along with the proportion of a binomial distribution, by the method of moments. The result is a weighted t-type test statistic.

### Generation of pET45b construct for His tag purification of *P. putida* KT2440 3-hydroxybutyrate dehydrogenase (hbdH) protein

Genomic DNA was isolated from 1 ml of *P. putida* KT2440 grown in LB at 30°C, shaken at 200 rpm for 16 h using a GeneJET genomic DNA purification kit (Thermo Scientific, Dublin, Ireland) according to the manufacturer’s instructions. The *hbdH* gene was amplified from the genomic DNA. The PCR product was gel purified from a 1% agarose gel using a GeneJET gel extraction kit (Thermo Scientific, Dublin, Ireland) and ligated into pGEM-T Easy vector (Promega, Madison, WI) using T4-DNA ligase. The ligation mixture was transformed in XL-10 gold competent cells (Stratagene, Agilent Technologies, Santa Clara, CA) by heat shock according to the manufacturer’s instructions. Colonies were tested by PCR for the presence of the hbdH gene. Positive colonies were inoculated into 2 ml of LB medium supplemented with 50 μg/l carbenicillin and grown for 16 h at 37°C, shaking at 200 rpm. Plasmid DNA was extracted from these cultures using a GeneJET plasmid miniprep kit (Thermo Scientific, Dublin, Ireland) and sent to GATC (Konstanz, Germany) for sequencing. The *hbdH* gene was excised from a sequence verified plasmid using restriction endonucleases (Promega, Madison, WI). The expression vector pET45b (Novagen, Madison, WI) was also digested with the same enzymes. The gene was ligated into the digested vector with T4 DNA ligase to generate the pET45b_hbdH expression vector. The ligation mixture was transformed into BL-21 Gold competent cells (Stratagene, Agilent Technologies, Santa Clara, CA) by heat-shock according to the manufacturer’s instructions. Positive transformants were selected by ampicillin resistance and confirmed by PCR.

### Expression of his-tagged hbdH in *E.coli* BL21 cells

*E. coli* BL21 cells containing the pET45b_hbdH plasmid were grown in 2 l shake flasks containing 400 ml of LB medium supplemented with 50 μg/ml carbenicillin at 25°C, shaking at 200 rpm until the OD600 reached 0.4 (approximately 6 hours). Cultures were cooled on ice for 30 minutes. Isopropyl β-D-1-thiogalactopyranoside (IPTG) was added to a final concentration of 0.5 mM and the cultures were grown at 25°C, shaking at 200 rpm for 18 h.

### Purification of His-tagged 3-hydroxybutyrate dehydrogenase protein

Cells were harvested by centrifugation at 3600 g for 12 minutes at 4°C. The supernatant was discarded, and cell pellets resuspended in lysis buffer (3ml Bugbuster mastermix (Merck Millipore, Cork, Ireland) and 1.5 ml binding buffer (300 mM NaCl, 20mM imidazole, 50 mM sodium phosphate) per 1 g of wet cell pellet. The resuspended pellets were incubated at 30°C for 30 minutes. Cell debris was removed by centrifuging for 30 minutes at 43,146g at 4°C. Cell lysate was filter using a sterile 0.45 μm filter. The cell lysate was then passed through a 1ml HisTrap column (GE healthcare, Little Chalfont, UK). For every 2 ml of cell lysate passed though the column, 2 ml of binding buffer was also passed through the column. The HisTrap column was then attached to an AKTA prime system (GE healthcare, Little Chalfont, UK).

The column was washed with 6ml binding buffer and then eluted using a gradient of elution buffer (300 mM NaCl, 500 mM imidazole, 50mM sodium phosphate). 2 ml fractions were collected. Fractions were analysed by SDS-PAGE under denaturing conditions. The resolving gel contained 12 % and stacking gel 4 % acrylamide (w/v). Fractions containing the hbdH protein were pooled and the protein concentration determined by BCA assay (21). 25 μl of each fraction to be measured was added in duplicate to a 96 well microtitre plate. 200 μl of a bicinchoninic acid solution containing 2 % (v/v) copper sulphate was added to each sample. The plate was incubated at 40°C for 30 minutes. The absorbance of each sample at 550 nm was measured using a SPECTROstar Nano microplate reader (BMG Labtech, Ortenberg, Germany).

### Assay for activity of 3-hydroxybutyrate dehydrogenase protein

The activity of the purified protein was measured using an assay previously described (22). NAD^+^, the co-factor for this enzyme is converted to NADH during the reaction. NADH can be measured spectrophotometrically at 340 nm. Assays were carried out in 200 μl volumes in a 96 well plate. 10mM substrate and 1.5 mM NAD^+^ were added to 50mM potassium phosphate buffer, pH 8. Enzyme was added to start the reaction, the plate was placed in a plate reader at 23°C and the change in absorbance at 340 nm was measured in a SPECTROstar Nano microplate reader (BMG Labtech, Ortenberg, Germany). Values were converted to NADH concentrations using an extinction coefficient of 6.3 mM^−1^cm^−1^. The natural substrate for the enzyme, 3-hydroxybutyrate was used as the positive control. Negative control reactions containing no enzyme, no NAD^+^ and boiled enzyme were also carried out.

## Results

### Generation of *P. putida* KT2440 Δ*glpK* deletion mutant

The *glpK* gene was successfully replaced with a gentamycin cassette on the chromosome to yield a *P. putida* KT2440 Δ*glpK* strain. The mutation was confirmed by gentamycin resistance screening, Southern blot technique and by DNA sequencing of the mutant chromosome.

### Growth of *P. putida* KT2440 wild type and Δglpk on glucose, sodium octanoate and glycerol

Growth of the wild type and Δglpk mutant was determined after 48h of cultivation in minimal salt medium containing 20 mM of the carbon source at 30°C and shaking at 200 rpm. All conditions were tested in a minimum of three separate shake flasks. Growth levels of the strains was comparable when grown on glucose or sodium octanoate as the sole carbon source (Figure 1). Minimal growth of the Δglpk mutant was detected when grown on glycerol for 48 hours (figure 1). When incubation was extended the Δglpk mutant achieved a final OD_540_ of 3.1 which is 1.6-fold lower than that observed for the wild type strain (Figure 2).

**Figure 1.**
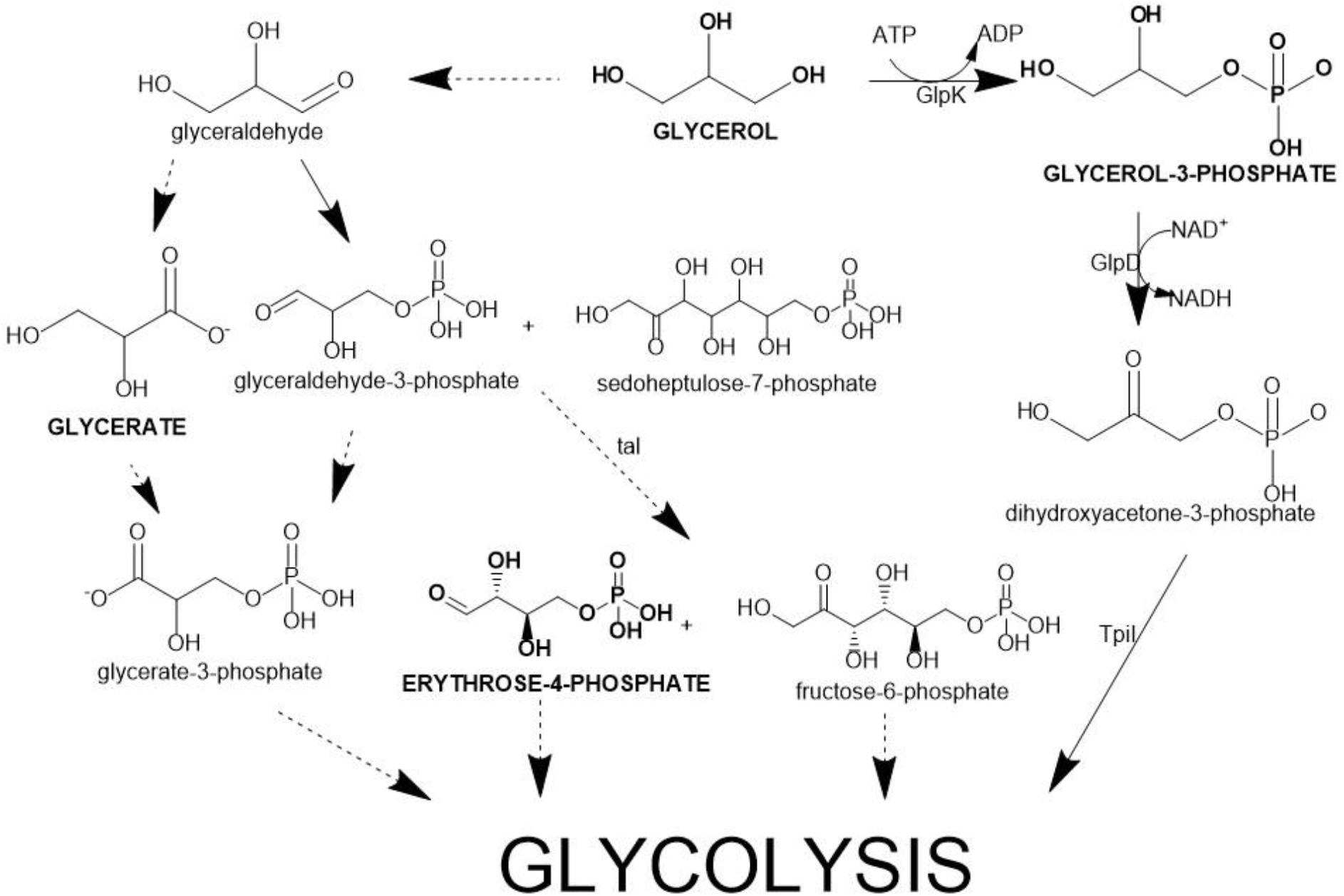
Glycerol metabolic pathway in *P. putida* KT2440 (solid arrows) and proposed metabolic pathway in the absence of glycerol kinase (dotted arrows). Metabolites detected at high levels (>1M counts) in the ΔglpK mutant are highlighted in bold and uppercase.

**Figure 2.**
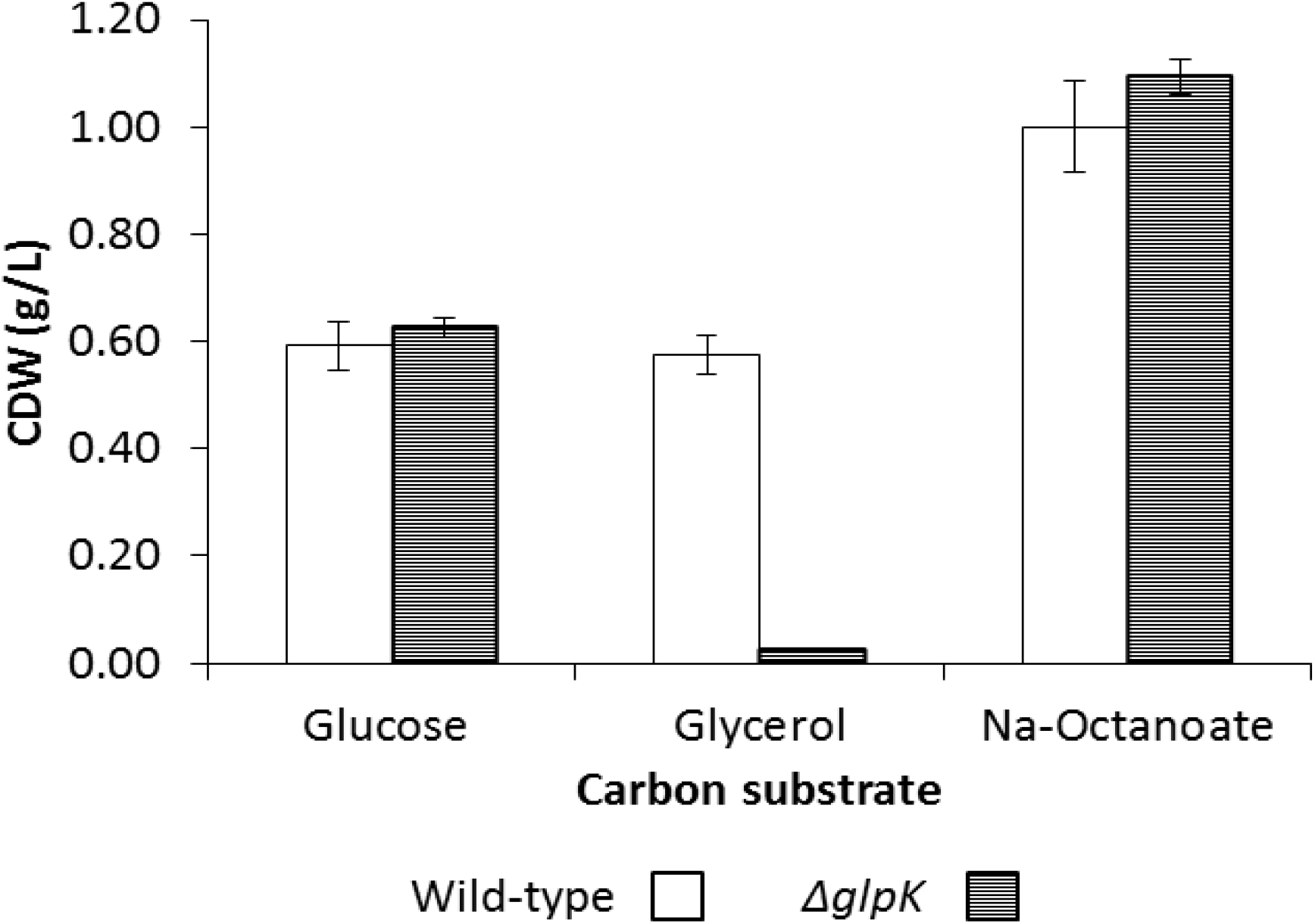
Growth of *P. putida* KT2440 and *glpK* on glucose, glycerol and sodium octanoate in minimal salts medium. Optical density (OD) was measured at 540 nm. When harvested? seems to not be in line with figure 3. Can figure 1 and 2 be presented as a and b.

### Complementation of *P. putida* KT2440 *ΔglpK* mutant

To show conclusively that the observed phenotype was as a direct result of the deletion of the gene and not by other polar effects caused by the mutagenesis process, a selectively inducible copy of the gene, harboured in the pJB8621 expression vector, was introduced into the mutant and wild-type strain and the ability of the strains to grown on glycerol was analysed.

The complemented mutant *P. putida* KT2440 *ΔglpK* pJB861/*glpK* reached near wild-type levels of biomass (96 % recovery). *P. putida* KT2440 *pJB861/glpK*, which is the wild type strain harbouring an induced extra copy of the *glpK* gene, grew to very similar levels to that of the wild-type strain (95 % of wild-type CDW) (Figure 3).

**Figure 3.**
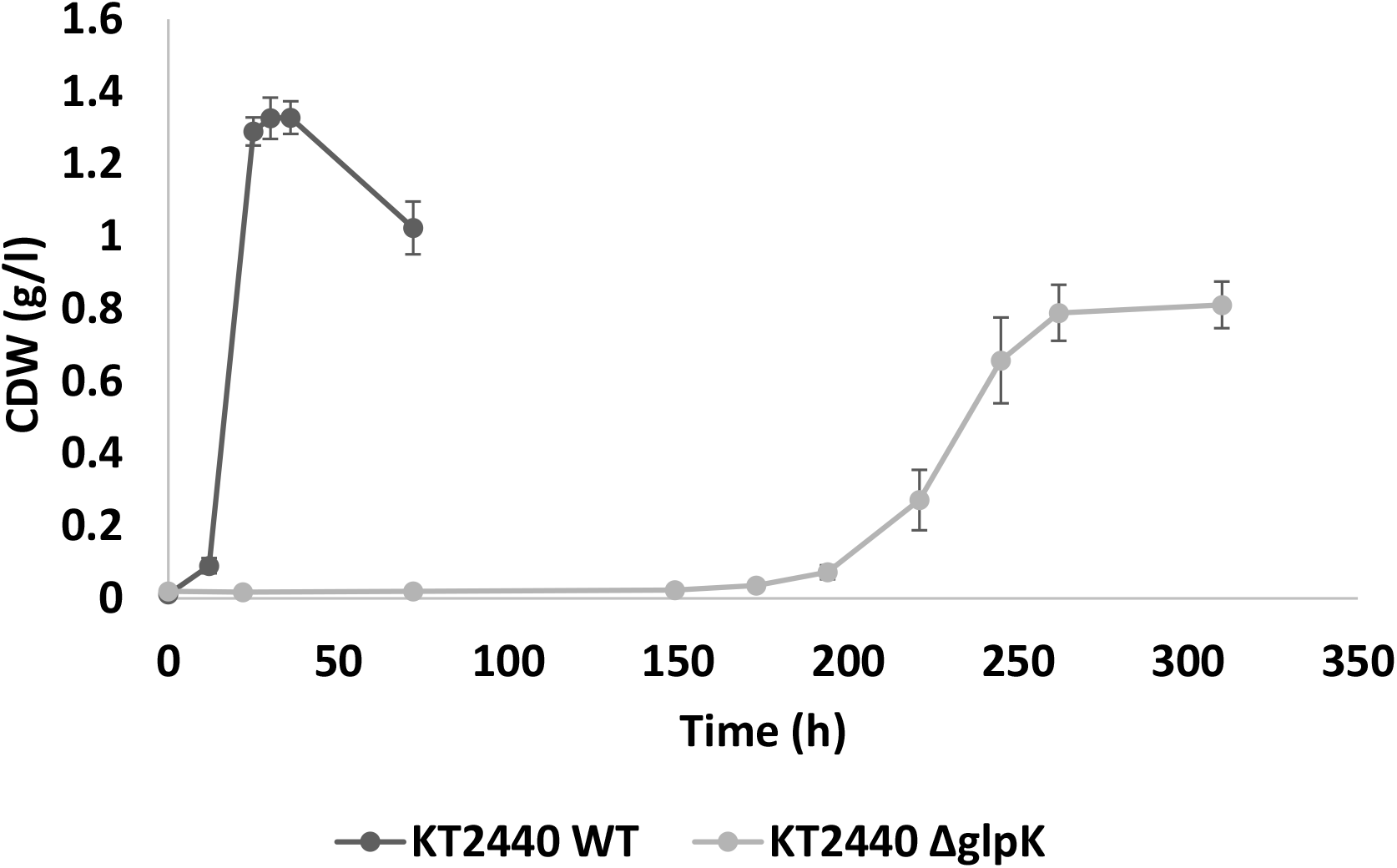
Growth of *P. putida* KT2440 wildtype and ΔglpK on glycerol as the sole carbon source. Optical density (OD) was measured at 540 nm.

### Transcriptomics of *P. putida* KT2440 wild type and ΔglpK grown on glycerol as the sole carbon source

RNA was isolated from *P. putida* KT2440 wild type and Δglpk cells in mid logarithmic phase of growth and it was sent to Baseclear (Leiden, Netherlands) for transcriptomic analysis. There were statistically significant differences in the expression levels of 2962 transcripts. 1757 of these are upregulated in Δglpk cells and 1205 are downregulated. Of these 2962 differentially expressed transcripts, 191 were more than 2-fold upregulated and 175 more than 2-fold down regulated.

The transcription of enzymes in the glycerol metabolic pathway were all down-regulated, ranging from 32-fold for glpK to 1.2-fold for glpR. As the *glpK* was knocked out by inserting a gentamicin resistance cassette in place of the gene, small parts of the gene remained on either side of the gentamicin resistance cassette. This resulted in a small number of short transcripts being assigned to glpK rather than 0 as may be expected. Other enzymes that are potentially involved in glycerol metabolism show small differences in transcription in wild type cells compared with ΔglpK cells. Metabolites from glycerol metabolism enter central metabolism through the glycolysis. All the enzymes in this pathway are slightly down regulated (1.2 − 1.6-fold) except for pyruvate kinase which is 1.5-fold upregulated. There were no significant differences in transcription of any genes involved in fatty acid β-oxidation or PHA synthesis. However, transcription of genes involved in short chain length fatty acid metabolism were highly upregulated in ΔglpK cells versus wild type cells (Table 1). 54 genes whose functions are unknown were more than 2-fold upregulated in the ΔglpK mutant compared to the wild type and 27 transcriptional regulators are also more than 2-fold upregulated in the ΔglpK mutant compared to the wild type. The change in expression of relevant genes is shown in Table 1.

**Table 1.**
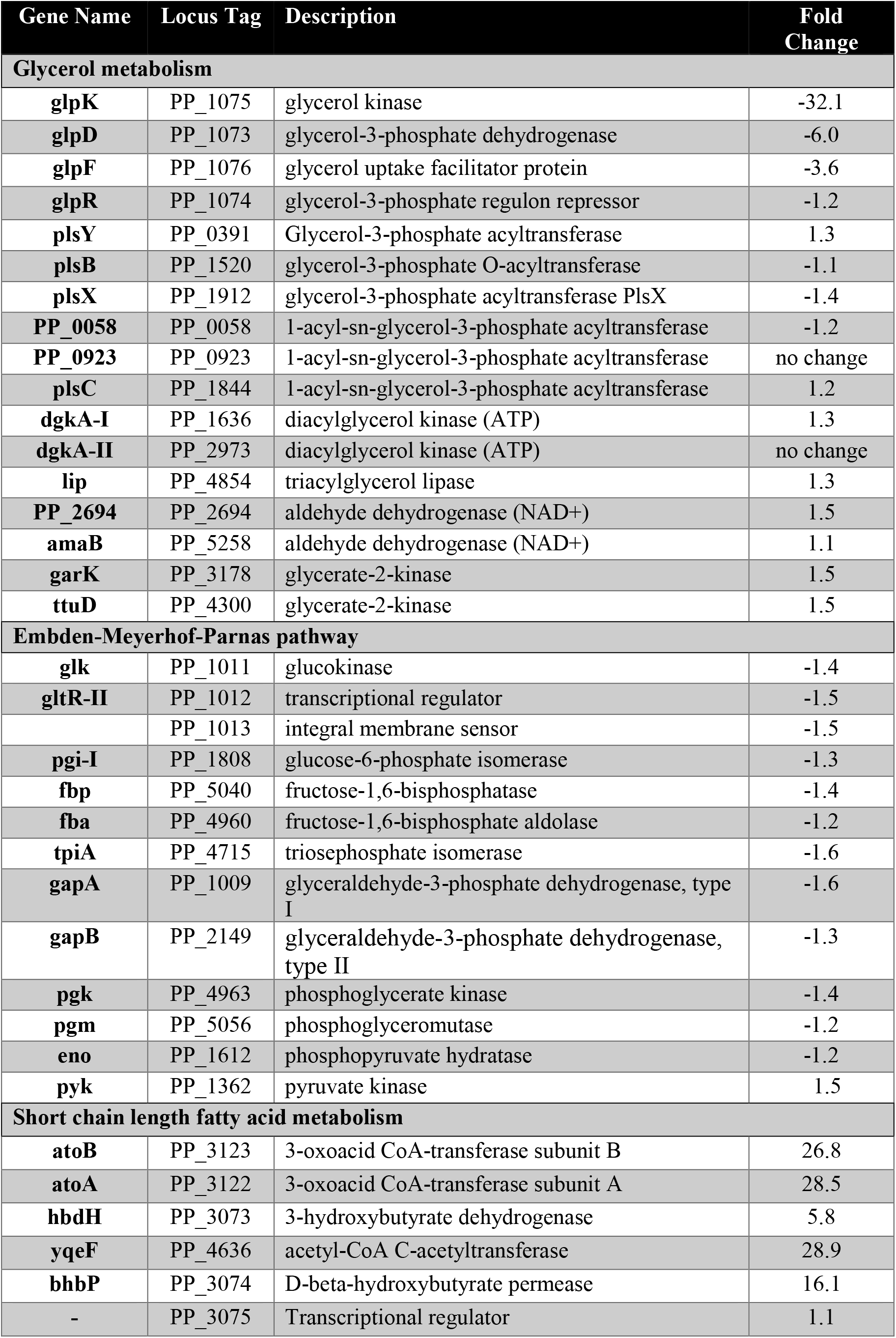
Change in expression of genes in ΔglpK versus wild type *P. putida* KT2440 cells in mid logarithmic phase grown on glycerol as the sole carbon and energy source.

### Metabolomic analysis of *P. putida* KT2440 wild type and ΔglpK cells grown on glycerol as the sole carbon source

Metabolites were extracted from *P. putida* KT2440 wild type and ΔglpK cells in mid logarithmic phase of growth. Metabolites potentially produced in glycerol catabolism as well as common metabolites from central metabolism were analysed in cell extracts and levels compared in wild type and ΔglpK cells (Figure 4). Production of glycerate is highly upregulated (21 fold) in the ΔglpK mutant. Furthermore erythrose-4-phosphate is detected in ΔglpK mutant cell extracts, but not in wild type cells. Interestingly levels of glyceraldehyde and 3-phosphoglycerate are similar in both strains. As expected, levels of glycerol-3-phosphate are much higher in the wild type strain, however some glycerol-3-phosphate is produced in the ΔglpK strain, suggesting that there may be a non-specific kinase acting on glycerol in the mutant strain. Dihydroxyacetone is detected only in the wild type strain. The upregulation of glycerate levels in cell extracts suggests that glycerol may be metabolised via this intermediate. This could be achieved through the action of a dehydrogenase, converting glycerol to glyceraldehyde and then to glycerate.

**Figure 4.**
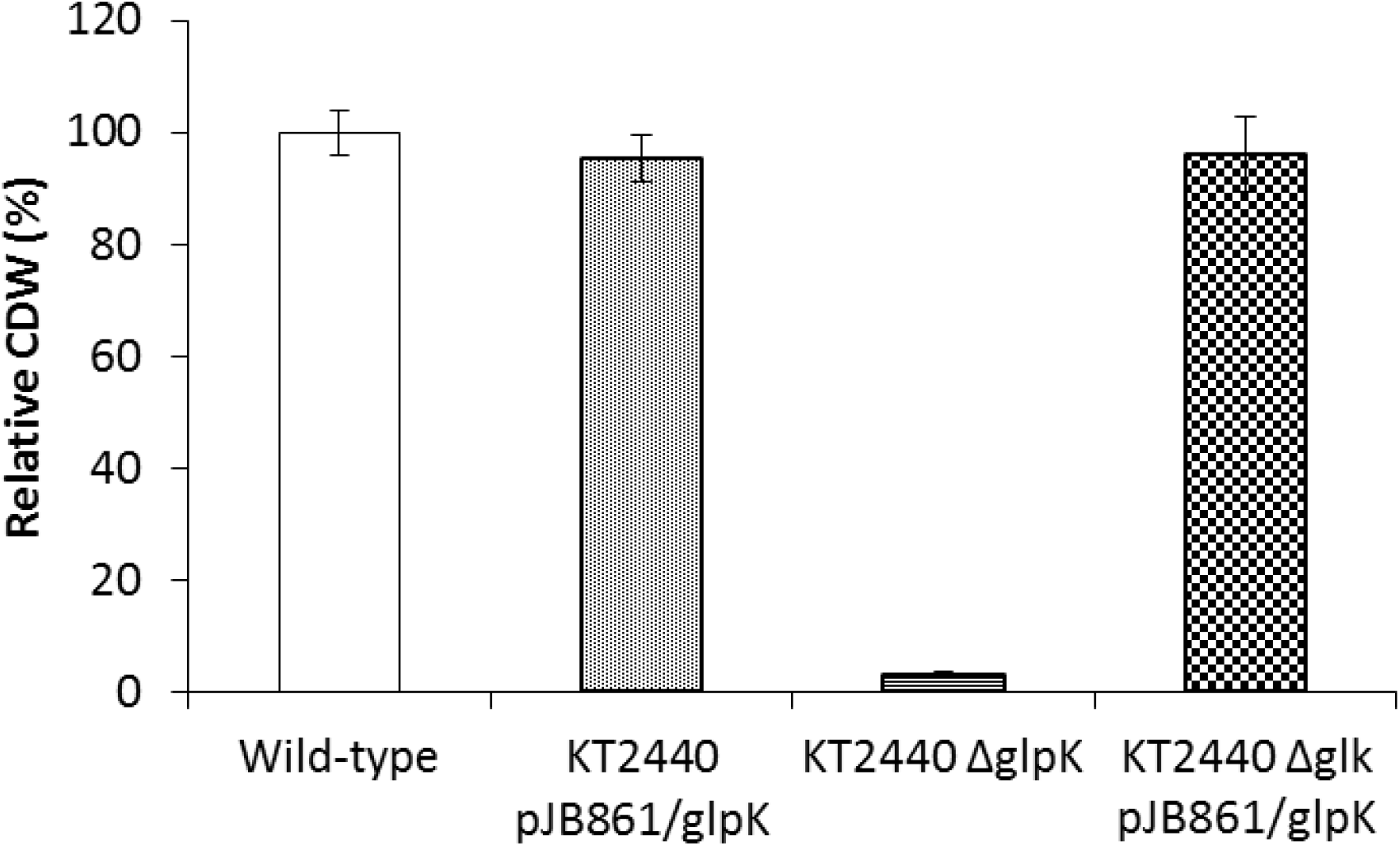
Growth of *P. putida* KT2440 wild type and ΔglpK cells, with a complemented copy of glpK relative to growth of the wild type *P. putida* KT2440. The glpk gene was expressed on pJB861 plasmid. Glycerol was supplied as the sole carbon and energy source.

### Cloning, expression and purification of 3-hydroxybutyrate dehydrogenase

Analysis of the transcriptome revealed that 3-hydroxybutyrate dehydrogenase (3HBDH) transcription was 5.8-fold up regulated in the ΔglpK mutant. To investigate if this enzyme could act on glycerol, the 770 bp gene was cloned into pET45b, which has an N-terminal 6 histidine tag.

The protein was expressed in *E. coli* BL21 cells and purified using a nickel affinity column on an Akta basic protein purification system. The protein was eluted from the column using a gradient of 500 mM imidazole. 2 ml fractions were collected, and the fractions run on a 12% SDS PAGE gel to determine which fractions contained the 3HBDH. Fractions containing the 3HBDH protein were pooled and assayed for activity. Protein concentration was determined by BCA assay.

### Assay for 3-hydroxybutyrate dehydrogenase activity

The purified protein was assayed for its activity towards 3-hydroxybutyrate, glycerol and glyceraldehyde. As NAD^+^ is a cofactor for the enzyme, NADH production was used as a measure of enzyme activity. Activity was detected when 3 hydroxybutyrate and glyceraldehyde were used as substrates. The rate of NADH production was much higher for 3-hydroxybutyrate than for glyceraldehyde (Figure 5). No production of NADH was observed when glycerol was used as the substrate. There was no production of NADH in any of the negative controls (using no enzyme, no NAD^+^ or enzyme that had been boiled at 100°C for 10 minutes).

**Figure 5.**
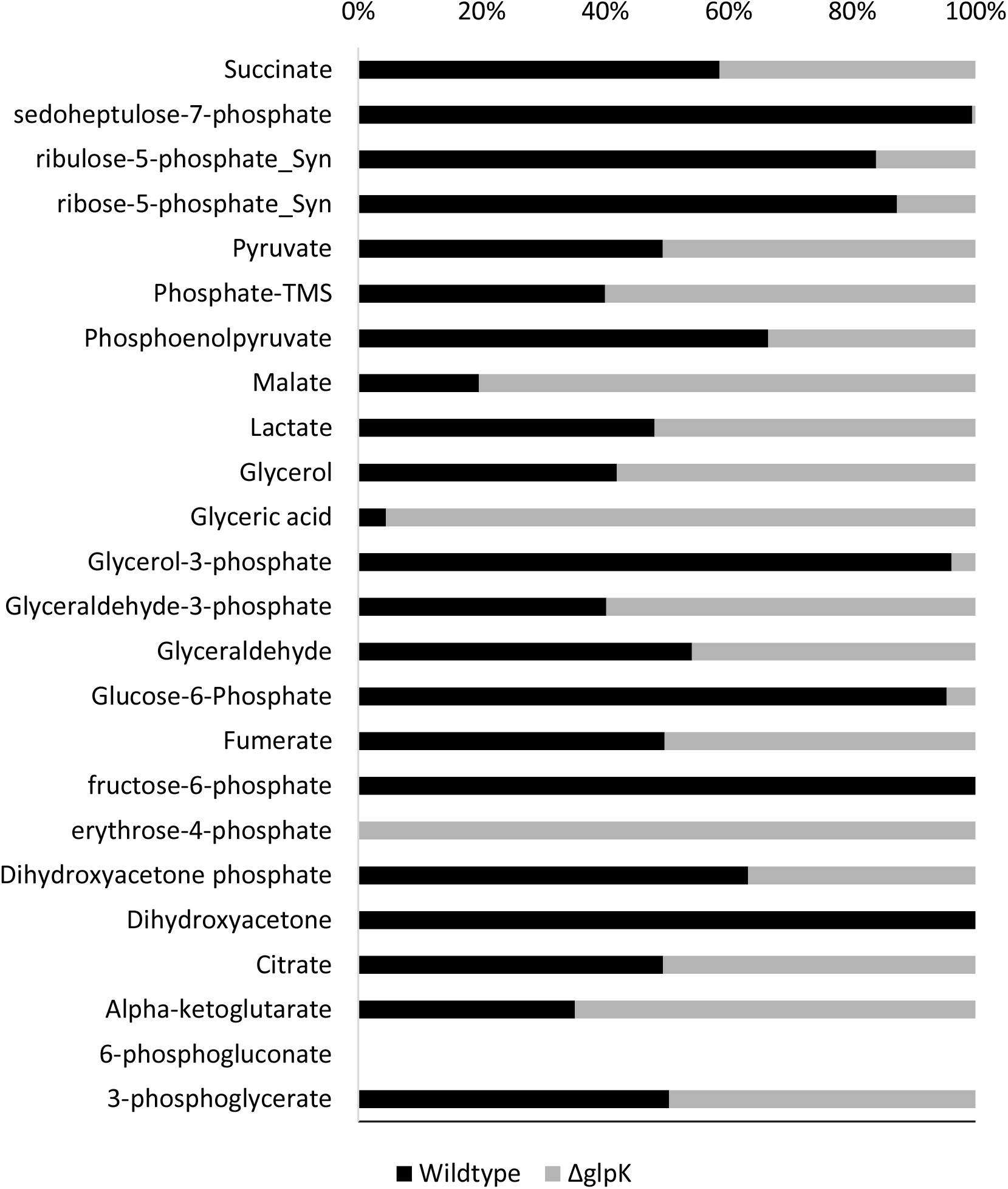
Metabolites identified in extracts of *P. putida* KT2440 wildtype and ΔglpK cells grown on glycerol as the sole carbon and energy source. Cells were harvested at the mid log phase of growth. Values expressed as a percentage of total metabolite detected in both samples.

**Figure 6.**
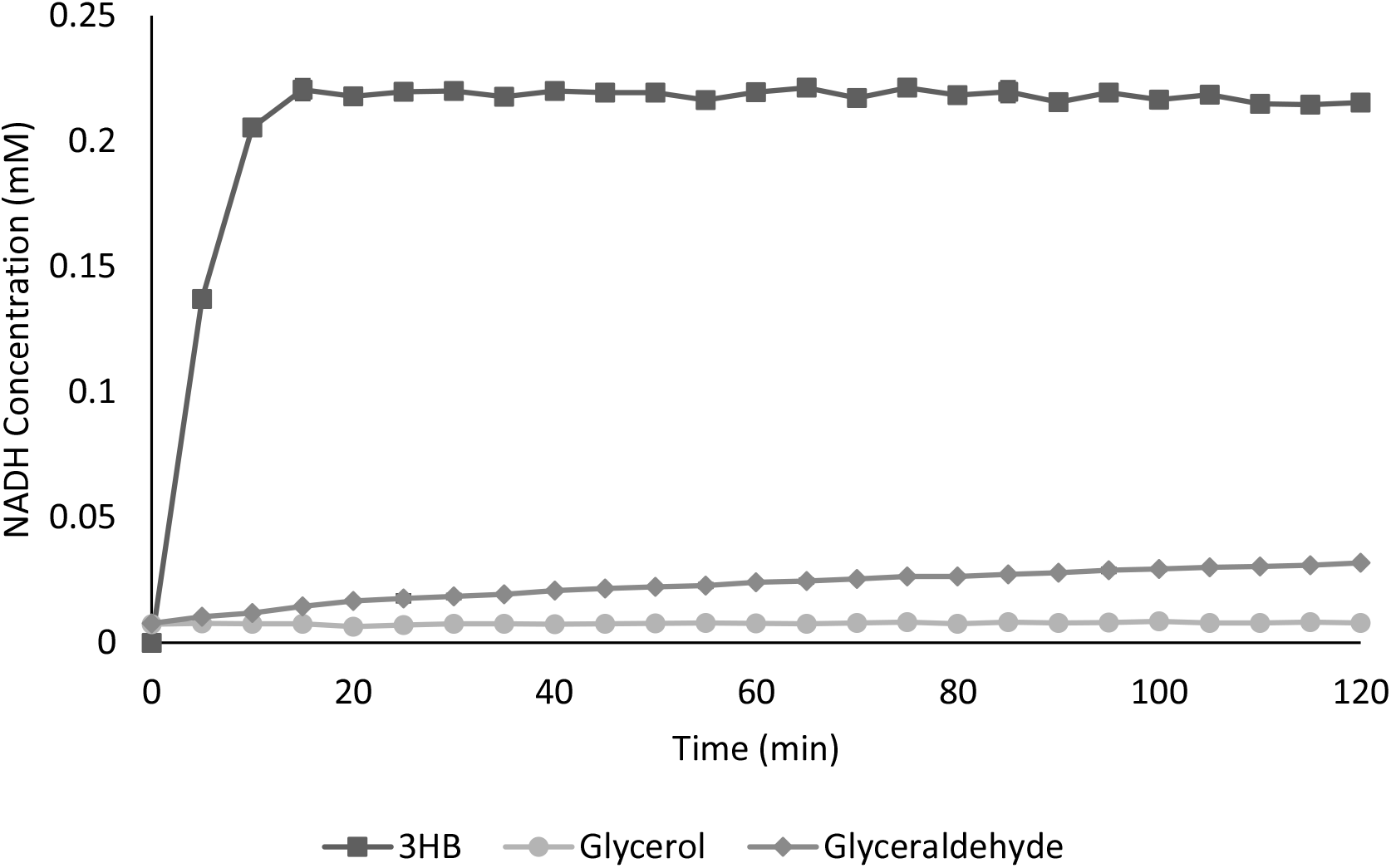
NADH production when 3 hydroxybutyrate (3HB), glycerol or glyceraldehyde was used as the substrate for recombinantly produced 3-hydroxybutyrate dehydrogenase from *P. putida* KT2440.

## Discussion

The first step in the proposed pathway for glycerol catabolism in *P. putida* KT2440 is phosphorylation by the glycerol kinase (PP_1075). We knocked out this gene and found that the knockout mutant retained the ability to grow on glycerol as the sole carbon source, though it had a much-extended lag period and achieved 1.6-fold less biomass than the wild type strain. The ΔglpK mutant grew similarly to the wild type when grown on glucose or sodium octanoate indicating the *glpK* gene is affecting early stage metabolism of glycerol and not affecting central metabolism (23). This is consistent with the literature where the *glpK* gene product is only induced in the presence of glycerol and is only be expressed at basal levels during incubation with glucose or sodium octanoate.(9,24)

It has previously been established in *E. coli* and *P. aeruginosa* that this *glpK* gene product is vital for the production of glycerol-3-phosphate and hence for the up-regulation of all major glycerol metabolic genes (1,25). However, we have found that the ΔglpK mutant still grows albeit with a lag period and a lower growth yield but an alternative glycerol metabolic pathway must exist for growth of *P. putida* KT2440 in the absence of the glycerol kinase.

In bacteria, there are two major pathways involved in glycerol metabolism. The phosphorylation pathway and the oxidation pathway (26). As previously outlined, *Pseudomonas putida* KT2440 uses the most common biological pathway namely the phosphorylation pathway. Other microorganisms such as *Klebsiella pneumoniae* use the oxidation pathway in which a glycerol dehydrogenase converts glycerol into dihydroxyacetone (27). Some facultative anaerobic bacteria such as *Klebsiella aerogenes* use the phosphorylation pathway under aerobic conditions, but can employ the oxidation pathway in the absence of oxygen (28). In some bacteria, it is also possible for glycerol to be oxidised to glyceraldehyde (29), which may then be converted to glycerate (30).

As expected, enzymes in the glycerol kinase pathway were down regulated in the ΔglpK mutant. However *glpR*, the regulator of expression of glycerol kinase is only 1.2-fold downregulated. This is in keeping with previous studies that show the levels of *glpR* remain the same regardless of carbon source used (31). There is a report of a glycerol kinase deletion mutant of *E. coli* K12 which could use glycerol as the sole carbon source for growth by using an NAD^+^ linked glycerol dehydrogenase to metabolise glycerol. That enzyme showed much higher activity towards dihydroxyacetone compared with glyceraldehyde (32). Metabolite analysis showed that no dihydroxyacetone was produced by our mutant strain when grown on glycerol, therefore this pathway is not likely to be employed for metabolism of glycerol in the absence of the glycerol kinase in *P. putida* KT2440.

Levels of glyceraldehyde and glycerol-3-phosphate are approximately the same in both wild type and mutant. However, levels of glycerate are highly upregulated in the mutant, suggesting that glycerol may be converted to glyceraldehyde and then to glycerate in the absence of the glycerol kinase. While transcription of glycerate kinase is 1.5-fold upregulated in the ΔglpK mutant strain compare to the wildtype, levels of 3-phospho-glycerate are similar in both strains.

Erythrose-4-phosphate is also detected in the mutant strain but not in the wildtype. Furthermore higher levels of glyceraldehyde-3-phosphate are also present in the mutant compared to the wild type KT2440. Transaldolase, the enzyme that converts glyceraldehyde-3-phosphate + sedoheptulose-7-phosphate to erythrose-4-phosphate + fructose-6-phosphate is 1.4-fold upregulated in the mutant. However, neither sedoheptulose-7-phosphate nor fructose-6-phosphate are detected in the mutant. As glycerate is upregulated in the mutant strain, it is possible that excess glyceraldehyde is also formed in the mutant strain, which may be easily converted to glyceraldehyde-3-phosphate, which can be converted to erythrose-4-phosphate. Glyceraldehyde-3-phosphate can enter the Embden-Meyerhof-Parnas pathway (33). Enzymes in this pathway are slightly downregulated in the mutant strain compared to the wildtype indicating that EMP is still used for glycerol metabolism by the ΔglpK mutant but the lower expression may contribute to the slower growth rate of the mutant compared to the wild type.

If glycerol is converted to glyceraldehyde in the mutant strain, there must be a dehydrogenase enzyme acting non-specifically in this strain. Transcription of the enzyme 3-hydroxybutyrate dehydrogenase was highly upregulated in the ΔglpK mutant compared to the wild type. 3-hydroxybutyrate dehydrogenase converts 3-hydroxybutyrate to acetoacetate in a reversible reaction with NAD^+^ as a cofactor. As it was upregulated in the mutant, it was hypothesized that it may also be able to non-specifically catalyse the conversion of glycerol to glyceraldehyde or glyceraldehyde to glycerate.

3-hydroxybutyrate dehydrogenase from *P. putida* KT2440 was expressed in *E. coli*, purified and tested for activity towards glyceraldehyde and glycerol. Activity was detected towards glyceraldehyde, albeit at a slower rate than towards 3-hydroxybutyrate, the natural substrate for the enzyme. However, no activity was detected towards glycerol. Analysis of the KT2440 genome revealed 5 further dehydrogenases that could be responsible for glycerate production in the mutant strain, these enzymes were between 2 and 2.7 fold upregulated in the transcriptomic analysis. A companion study to this work, which investigated the physiological responses of *P. putida* KT2440 towards rare earth elements during growth with different growth substrates, identified that two periplasmic PQQ-dependent alcohol dehydrogenases that are essential to initiate an alternative glycerol pathway via the oxidation of glycerol to glyceraldehyde (34). No significant difference in the expression of these PQQ-dependent dehydrogenases was observed in the transcriptomic analysis. Now that an alternative glycerol pathway in *P. putida* KT2440 has been discovered it opens up the possibilities for studies into the regulation of both pathways in which could increase the efficiency of glycerol consumption in this strain and open up new biotechnological possibilities.

In conclusion a glycerol kinase negative mutant of *P. putida* KT2440 is capable of growth using glycerol as a sole carbon and energy source with a very long lag phase. Transcriptome and metabolome analysis suggest glycerate and erythrose-4-phosphate as major metabolites in the mutant strain. Glyceraldehyde and glyceraldehyde-3-phosphate are also present in the mutant but at similar levels to the wild type strain. Short chain fatty acid metabolism genes are upregulated in the Δglpk mutant. One of these enzymes, 3-hydroxybutyrate dehydrogenase, has activity towards glyceraldehyde which could explain the increase in the concentration of glycerate in the ΔglpK mutant.

### Funding information

Meg Walsh was funded by the Irish Research Council Employment Based Postgraduate Programme (Ref no. 39826). The authors acknowledge support under EU Horizon 2020 research and innovation programme 633962 for the project P4SB. LMB acknowledges also funding by the Cluster of Excellence “The Fuel Science Center - Adaptive Conversion Systems for Renewable Energy and Carbon Sources”, which is funded by the Excellence Initiative of the German federal and state governments to promote science and research at German universities.

